# Complexity of the Relationship between Life Expectancy and Overlap of Lifespans

**DOI:** 10.1101/195867

**Authors:** Julia A. Barthold, Adam Lenart, Annette Baudisch

**Affiliations:** Max-Planck Odense Center on the Biodemography of Aging, University of Southern Denmark, Odense, Denmark; Department of Public Health, University of Southern Denmark, Odense, Denmark; Department of Biology, University of Southern Denmark, Odense, Denmark

## Abstract

Longevity has long been recognised as a key facilitator of reciprocal altruism because repeated cooperation of partners hinges on mutual survival. Although demographic tools can be used to quantify mutual survival and expected overlapping lifespans, studies on the evolutionary theory of cooperation take only limited advantage of demography. Overlap of lifespans depends on variation in survival across ages and can be high or low independently of high or low life expectancies. Here we develop formal demographic measures to study the complex relationships between shared life expectancy of two birth cohort peers, the proportion of their lives that they can expect to overlap, and longevity. We simulate age-specific mortality schedules using a Siler model to reveal how infant and senescent mortality, along with age-independent mortality, affect the relationship between the proportion of life shared and life expectancy. We find that while the proportion of life shared can vary vastly for similar life expectancies, almost all changes to mortality schedules that result in higher life expectancies also result in higher proportions of life shared. A distinct exception occurs if life expectancy increases due to lowering the rate of senescence. In this case the proportion of life shared decreases. Our work shows that almost all selective pressures that result in higher life expectancies also result in a larger proportion of life shared. Therefore, selective forces that extend life also improve the chances that a cooperative system would be stable in terms of reciprocal interactions. Since reciprocal interactions may also reduce mortality and result in a feedback loop with the evolution of longevity, our measures and findings can be used for future cross-species comparisons that aim to disentangle predecessor and successor in the evolution of longevity and cooperation.

## Introduction

Many primate species engage in unidirectional or reciprocal cooperation with others [1, 2]. Dyadic interactions with relatives or other individuals are particularly common among humans. This cooperative behaviour has presumably evolved because it increases the fitness of the individual who performs the behaviour by yielding either indirect or direct fitness gains [3]. Direct fitness gains via reciprocity are often not immediate but rather occur with a time delay to be realised in a future interaction between the cooperation partners [4, 5]. The probability that two individuals will repeatedly interact is therefore integral to theoretical models of reciprocal cooperation [6]. This probability depends among other factors on the survival of both interaction partners [7]. Despite the central role of mutual survival in determining the uncertainty of a future reciprocal interaction, age-structured demography has only marginally been integrated into the theory of the evolution of cooperation and only as part of individual-based models [e.g. 8]. As a first step towards this aim, we study here mutual survival of individuals using stable population theory.

A human baby born today in an industrialised country can expect to share most of its lifetime with a peer from its birth cohort due to high lifespan equality (i.e. most individuals live similarly long) [9]. High lifespan equality arises from a rectangularised survival function, which captures the fact that most individuals will survive to a similar age [10]. What is true for humans today, however, need not be true in general. Across the tree of life, species show an astounding diversity of survival functions, with remarkable differences even between human populations [11]. In this work we ask how different survival functions determine, firstly, life expectancy, secondly, the expectancy of overlapping life among two peers of a birth cohort (”shared life expectancy”), and thirdly, the proportion of shared life expectancy in relation to life expectancy (”proportion of life shared”). A low proportion of life shared adds uncertainty to the future availability of reciprocal cooperation partners and thus may hinder the evolution of cooperation.

Species with similar life expectancies can vastly differ in the shape of their survival function and its underlying age-specific mortality rates. In species characterised by falling mortality a large proportion of individuals die during infancy due to high infant mortality, but falling mortality rates with age guarantee a long life for the survivors. On the other hand, rising morality indicates that individuals tend to survive childhood, but then mortality rates increase with age. Furthermore, mortality rates can be constant across all ages (“age-independent mortality”) or result in “bathtub mortality” with high infant mortality, low mortality during adult ages, followed by an increase in mortality at senescent ages. Any of these mortality trajectories can result in similar life expectancies from birth. Methods have been developed to distinguish these two aspects of the ageing regime as the “pace and shape” of ageing [12–14], where life expectancy is a measure of pace and the relative increase or decrease in mortality across the life course is a measure of shape.

Although we know that similar life expectancies can result from a range of mortality shapes, it is not clear how shared life expectancy and the proportion of life shared relate to these different mortality shapes. If most individuals die young, does this translate into two individuals being certain to overlap most of their short lives, or will the proportion of life shared be small because some individuals die young whereas others are lucky to survive infancy and reach advanced ages? Similarly, if all individuals survive infancy, but then mortality increases with age, does that mean that individuals at birth can expect to spend most of their lives overlapping with an individual born at the same time? And how does this change if mortality goes up quickly with age or only moderately?

To answer these questions, we first suggest formal demographic measures for shared life expectancy and for the proportion of life shared, based on stable-population theory. We then explore the relationships between life expectancy and the proportion of life shared using a parametric mortality model to simulate different mortality shapes. Mortality is modelled with three components: infant mortality, senescent mortality, and age-independent mortality. Most species for which age-specific mortality rates are known show a composite of these three components, often resulting in a bathtub shape of mortality (e.g. in mammals). The Siler model [22] is a convenient parametric model that captures these age-specific mortality rates. The model has proven a useful tool in ageing research, not least because each of its five parameters have a distinct biological interpretation, describing the different mortality components (see section: the parametric mortality model). Our aim is to identify how the different mortality components affect the relationship between life expectancy and the proportion of life shared, to be able to interpret empirical findings in future work, and to answer the question of whether increasing life expectancies are inevitably associated with an increasing proportion of life shared, or under which scenarios life expectancy can increase without the proportion of life shared following suit.

## Materials and Methods

### The parametric mortality model

The Siler model captures a competing risk hazard function composed of three summands that each represent one of the different components of mortality: infant mortality, age-independent mortality, and senescent mortality [22]. Given age *x* and a vector of mortality parameters ***θ***^T^ = [*a*_0_*, a*_1_*, c, b*_0_*, b*_1_], the model has the form

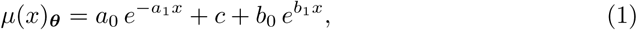

where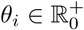

The first summand models the decrease in mortality rates that typically characterises infant ages, with *a*_0_ being the initial level and *a*_1_ modelling the rate of decrease. The middle summand captures age-independent mortality in the form of a constant hazard *c*, which is also known as the Makeham term [23]. The last summand is the Gompertz law of mortality [24], which captures the exponential increase in mortality rates with age from an initial level *b*_0_ with a rate of increase of *b*_1_.

While the hazard function of the Siler model facilitates the biological interpretation of the parameters, we used the survival function of the Siler model for calculations. It describes the probability to survive from birth to age *x* and is for any vector of mortality parameters ***θ*** given by

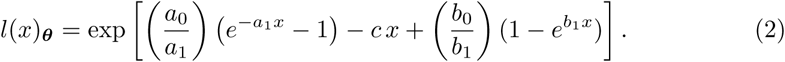

### Life expectancy, shared life expectancy, and the proportion of life shared

Based on the life expectancy at birth defined by 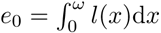we propose to measure the shared life expectancy at birth as 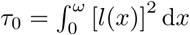This describes the mean number of years that two peers of a birth cohort will both be alive. Its upper bound is the life expectancy.

The proportion of life shared, i.e. the proportion of an individual’s life that she can expect to overlap with a birth cohort peer, is then given by

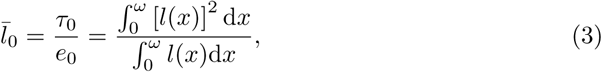

with the radix *l*(0) = 1. Incidentally, after developing the measure we found 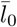 equals 1 – *G*_0_, where *G*_0_ is the Gini coefficient, which is commonly used to measure lifespan equality [14, 25]. This illustrates the close positive association between the proportion of life shared and lifespan equality.

### Mortality trajectory simulations

To study the relationships between *e*_0_, *τ*_0_, and 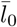 when the shape of mortality varies, we generated mortality trajectories by first producing sequences of all five mortality parameters, ranging from a minimum to a maximum value. The parameters ranges, and sequence lengths in brackets, where as follows: 0 *≤ a*_0_ *≤* 1.5 (10); 0 *≤ a*_1_ *≤* 1 (10); 0 *≤ c ≤* 0.015 (7); 0 *≤ b*_0_ *≤* 0.02 (30); and 0 *≤ b*_1_ *≤* 0.5 (30). All sequences had equal step lengths, apart of the *c* sequence that had one initial step of 0.005, followed by a step length of 0.002.

All permutations of the sequence elements together formed the range of mortality trajectories that we evaluated. We excluded all mortality trajectories for which the integrals in Eq (3) were not closed after 1000 years (5605 out of 630000). The upper integration limit in Eq (3) was set to the highest age at which at least 0.01% of a synthetic cohort was still alive. All analyses were conducted using the statistical programming language R version 3.1.3 [26]. Integrals were computed using R’s integrate()-function.

To understand the impact of the different components of mortality on the relationship between life expectancy and the proportion of life shared, we systematically subset the simulated mortality trajectories for ease of interpretation of results. We first turn to those mortality trajectories, where mortality rates decrease over age (falling mortality), then to those, where mortality increases with age (rising mortality), and thereafter to bathtub-shaped mortality rates (bathtub mortality).

## Results

### Falling mortality

Species that experience falling mortality are those characterised by mortality trajectories where mortality rates are high during infancy and decline with age. The parameters of interest are *a*_0_ and *a*_1_, which are the level and rate of decrease of infant mortality, respectively. We further modulate the level of age-independent mortality, governed by the parameter *c*, without which some mortality trajectories could result in immortal individuals. Infant mortality rates at each age are determined by the interaction of the level and rate parameter. For example, infant mortality can be low because both the level and rate of decrease are high or because the level is very low at birth, in which case the rate of decrease plays a less important role.

To illustrate the effect of infant mortality on the relationship between life expectancy and the proportion of life shared, Fig 1 depicts all mortality trajectories with the parameters *a*_0_, *a*_1_, and *c >* 0 and the senescent mortality parameters *b*_0_ = *b*_1_ = 0. We coloured the data points according to the ratio of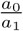, in order to illustrate the effect of infant mortality as a whole (i.e. to capture the interaction between level and rate).

Fig 1, panel a, reveals a J-shaped relationship between 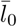 and *e*_0_. Values of 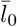 are highest for the shortest and longest lifespans and hit a minimum at intermediate lengths of life. The proportions of life shared are high when life expectancies are low because high levels of infant mortality cut all lives equally short. Similarly, the proportions of life shared are high when life expectancies are high because low levels of infant mortality allow all lives to be long. At intermediate levels of infant mortality, some lives are short while others are long, hence the low point in the proportions of life shared.

Fig 1, panel b, demonstrate the effect of the age-independent mortality level *c*. While it has little effect if levels of infant mortality dominate, *c* alters the slope of the positive relationship between 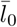 and *e*_0_ when infant mortality is lower. If *c* is low, then higher levels of *e*_0_ are reached for similar values of 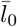, i.e. the slope increases with increasing levels of *c*. Therefore, similar values of age-independent mortality *c* can be associated with any degree of the proportion of life shared.

**Fig 1. The influence of infant mortality on the relationship between** 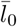 **and** *e*_0_ **(in years)**

Shown are data points calculated from mortality trajectories where *a*_0_, *a*_1_, and *c >* 0, and *b*_0_ = *b*_1_ = 0. (a) The effect of infant mortality. (b) The effect of age-independent mortality.

### Rising mortality

In the rising mortality setting we focus on mortality trajectories with zero infant mortality and rising mortality rates over age. The parameters of interest are *b*_0_ and *b*_1_, which are the level and rate of the increase in senescent mortality, respectively.

Furthermore, we also investigate the joint effect of senescent mortality and age-independent mortality, which is governed by the parameter *c*.

Generally, decreasing the level of senescent mortality increases both 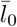 and *e*_0_ (Fig 2, panel a). That means that individuals both live longer, since the onset of senescent mortality is postponed, and overlap a larger proportion of their life with birth cohort peers. This observation holds with one exception. Where the age-independent mortality *c* is high, the relationship between 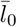 and *e*_0_ with decreasing levels of senescent mortality is hump-shaped (Fig 2, panel a and c). The negative relationship for lower *b*_0_ at high *c* values arises because under these circumstances the probability density function of age at death has a very long tail. The effect of this is to increase *e*_0_ but decrease 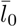, because a few individuals live to very old age.

Contrary to the effect of decreasing the level of senescence, decreasing the rate of senescence (*b*_1_) results in a negative relationship between 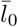 and *e*_0_ (Fig 2, panel b). This arises because decreasing the rate of senescence results in a progressively longer tail of the probability density function of age at death. The slope of the relationship depends on the level of senescence as it becomes more negative with increasing *b*_0_ values.

**Fig 2. The influence of senescent mortality, mediated by age-independent mortality, on the relationship between** 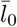 **and** *e*_0_ **(in years)**

Shown are data points calculated from mortality trajectories where *b*_0_ and *b*_1_ *>* 0, *c ≥* 0, and *b*_0_ = *b*_1_ = 0. (a) The effect of the level of senescent mortality (*b*_0_). (b) The effect of the rate of senescent mortality (*b*_1_). (c) The effect of age-independent mortality *c*.

### Bathtub mortality and age-independent mortality

Our results for bathtub mortality resemble the results for falling and rising mortality. For this part of the analysis, we first fixed infant mortality at a given level and let *b*_0_ and *b*_1_ vary. Similar to our findings for rising mortality, decreasing *b*_0_ results in a positive relationship between 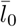 and *e*_0_ (Fig 3, panel a), while decreasing *b*_1_ results in a negative relationship between 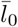 and *e*_0_ (Fig 3, panel b).

We then fixed the senescent part of the mortality trajectory and let the infant mortality parameters vary (Fig 4). The interaction between *a*_0_ and *a*_1_ in shaping infant mortality makes these patterns harder to discern. At high levels of *a*_0_, varying *a*_1_ results in a J-shaped relationship between 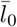 and *e*_0_. Decreasing levels of *a*_0_ attenuate the J-shape become until the relationship between 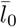 and *e*_0_ approaches a simple positive shape at the lowest values of *a*_0_ (*a*_0_ = 0.333). At the lower end of this positive relationship more individuals die during infancy then at the upper end of the relationship, where due to a high *a*_1_ value most individuals will survive infancy, which increases both 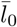 and *e*_0_, hence the positive relationship. Finally, in S1 Appendix we show that if mortality is constant over all ages, then 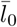 takes on a constant level and equals 0.5 for all life expectancies.

**Fig 3. The influence of senescent mortality on the relationship between** 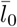 **and** *e*_0_ **(in years) when overall mortality rates are bathtub-shaped**

Shown are data points calculated from mortality trajectories where *a*_0_ = 0.5, *a*_1_ = 0.778, and *c* = 0. (a) The effect of the level of senescent mortality (*b*_0_). (b) The effect of the rate of senescent mortality (*b*_1_).

**Fig 4. The influence of infant mortality on the relationship between** 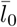 **and** *e*_0_ **(in years) when overall mortality rates are bathtub-shaped**

Shown are data points calculated from mortality trajectories where *b*_0_ = 0.0021, *b*_1_ = 0.121, and *c* = 0. (a) The effect of the level of senescent mortality (*b*_0_). (b) The effect of the rate of senescent mortality (*b*_1_).

## Discussion

In this study we have shown how different mortality components affect the relationship between the proportion of life shared and life expectancy. Most changes to age-specific mortality trajectories result in an increase of both measures and therefore in a positive relationship. The only circumstance where life expectancy increases and the proportion of life shared always decreases is when life expectancy increases due to lowering the rate of senescence (*b*_1_). While alterations to the age-specific mortality trajectories generally result in a positive relationship for most of the range of life expectancies, the relationship can also be negative over some life expectancies and J-shaped or hump-shaped over the whole range. This holds particularly for the case where infant mortality drops from extremely high values to lower values, which we have both observed for the falling mortality and the bathtub mortality setting. Finally, lowering the level of senescence (*b*_0_), which generally results in a positive relationship between the proportion of life shared and life expectancy, can result in a negative relationship over some values of life expectancy, if age-independent mortality is high. It is important to note that while we treat age in our analysis as starting from birth, the interpretation of our results holds similarly for treating age as starting from maturity, in which case the falling mortality setting corresponds to ”negative senescence” defined as a decline in mortality with age after reproductive maturity [27]–a phenomenon ascribed to prolonged and sustained development, growth, and regeneration.

According to life history theory, age-specific mortality rates are under strong selection [28]. Cross-species patterns of mortality therefore reflect species-specific adaptations to the environment, where energy expenditure into growth, survival, and fertility are traded-off against each other to maximise fitness [29]. High life expectancies are characteristic for slow life histories, where individuals grow to larger body sizes, invest heavily in survival, and give birth to few offspring repeatedly over the life course. From our results we can infer that selection pressures which increase life expectancy almost always increase the proportion of life shared, or in other words lifespan equality, and vice versa. A co-occurrence of both carries therefore little indication as to whether high proportions of life shared may aid the evolution of high life expectancies through enhancing cooperative behaviour, or whether high life expectancies inevitably co-occur with high proportion of life shared, which then may be a precondition for the evolution of cooperation.

Despite the fact that Axelrod and Hamilton [7] identified longevity as a key facilitators of reciprocal altruism, the literature that includes survival into the theory of cooperation is almost exclusively focussed on cooperation among relatives and on cooperative breeding [17, 19, 20]. A recent analysis of the co-evolution of longevity and cooperative breeding in birds showed that longevity preceded the evolution of cooperative breeding [18]. Theoretical work supports the notion that longevity promotes the evolution of cooperation [21]. Nevertheless, assuming the opposite causality, namely an effect of different modes of cooperative behaviour on the evolution of age-specific mortality and thus life expectancy, remains a tempting idea, and is also supported by theoretical models [8, 16]. Sociality among vertebrates is both common and patchily distributed, which points towards multiple evolutionary origins and consequently to the potential for multiple evolutionary advantages [30]. Therefore, while longevity seemed to have preceded the evolution of cooperative breeding, this may not hold true for other modes of cooperation such as cooperative territory defence or hunting.

The rise in human record life expectancy over the last 150 years may provide further insights [31]. The doubling of life expectancy seen during this period is one of the most remarkable examples for phenotypic plasticity in a trait under strong selection [32]. Initial gains in life expectancy came from reductions in infant mortality and young adult mortality, whereas since the 1950s progress has been driven by survival improvement at older ages [33]. Compelling evidence shows that survival improvements among the elderly has been propelled by the onset of senescent mortality being postponed [34]. This is equivalent to the level of senescent mortality decreasing rather than the rate of senescence falling. We have shown that gains in life expectancy through either bringing down infant mortality or decreasing the level of senescent mortality inevitably result in an increase in the proportion of life shared. The exceptional availability of data on humans may provide an opportunity to investigate the interplay between the extraordinary sociality of humans and the rise of prolonged, overlapping lifespans.

The relationship between life expectancy and overlapping lifespans already intrigues human demographers, who often express lifespan equality as lifespan disparity to illustrate a rising divide between countries or socio-economic classes within countries despite a general increase in life expectancy [35, 36]. Understanding how variation in lifespans arises from the shape of the mortality curve over age is therefore a research question that brings together disparate fields such as evolutionary biology, ageing research, economy, and demography. These diverse disciplines use different scale-free shape measures and interpret them as lifespan equality, lifespan disparity, or as we do here, as proportion of life shared. The cross-disciplinary interest in the relationship lends further weight to the importance of this research topic.

## Conclusion

According to life history theory, age-specific mortality rates are under strong selection. Almost all changes to age-specific mortality rates that result in higher life expectancies either through plasticity (as observed in humans) or selection (as observed cross-species) also result in a larger proportion of life shared. Individuals therefore live longer, and overlap longer, with individuals of their own and adjacent birth cohorts. Therefore, selective forces that extend life also improve the chances that a cooperative system would be stable in terms of direct or network-mediated reciprocal interactions. sReciprocal interactions may also reduce mortality and result in a feedback loop with the evolution of cooperation. Cross-species comparisons, co-evolution models, and the development of formal demographic measures for overlapping lifespans will in future work allow to disentangle predecessor and successor, cause and effect.

## Supporting Information

**S1 Appendix. Proof that** 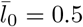 **when mortality is constant across ages.**

## Acknowledgments

We thank Jim Vaupel for suggesting the measure of 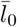 and initiating the line of research. We further thank Owen Jones for helpful comments on a previous version of the manuscript.

